# Severe neural tube defects due to failure of closure initiation can arise without abnormality of neuroepithelial convergent extension

**DOI:** 10.1101/2021.07.04.451044

**Authors:** Oleksandr Nychyk, Gabriel L. Galea, Matteo Molè, Dawn Savery, Nicholas D.E. Greene, Philip Stanier, Andrew J. Copp

## Abstract

Planar cell polarity (PCP) signalling is vital for initiation of neural tube closure in mice, with diminished convergent extension (CE) cell movements leading to a severe form of neural tube defect (NTD), termed craniorachischisis (CRN). Some human NTDs are also associated with PCP gene mutations, but affected individuals are generally heterozygous, whereas PCP homozygosity or compound heterozygosity is needed to produce CRN in mice. This suggests human NTDs may involve other genetic or environmental factors, that interact with partial loss of PCP function. We found that reduced sulfation OF glycosaminoglycans (GAGs) interacts with heterozygosity for the *Lp* allele of *Vangl2* (a core PCP gene), to cause CRN in mice. Here, we hypothesised that this GAG-PCP interaction may regulate convergent extension movements, and hence lead to severe NTDs in the context of only partial loss of PCP function. Both *Lp* and null alleles of *Vangl2* gave similar findings. Culture of E8.5 embryos in the presence of chlorate (a GAG sulfation inhibitor), or enzymatic cleavage of GAG chains, led to failure of NT closure initiation in the majority of *Lp/+* embryos, whereas few *+/+* littermates exhibited CRN. The chlorate effect was rescued by exogenous sulphate. Surprisingly, DiO labeling of the embryonic node demonstrated no abnormality of midline axial extension in chlorate-treated *Lp/+* embryos that developed CRN. In contrast, positive control *Lp/Lp* embryos displayed severe convergent extension defects in this assay. Morphometric analysis of the closure initiation site revealed abnormalities in the size and shape of somites that flank the closing neural tube in chlorate-treated *Lp/+* embryos. We conclude that severe NTDs involving failure of closure initiation can arise by a mechanism other than faulty neuroepithelial convergent extension. Matrix-mediated expansion of somites, flanking the closing neural tube, may be required for closure initiation.

## INTRODUCTION

Neurulation is the series of embryonic events that gives rise to the closed neural tube (NT), the precursor of the brain and spinal cord. Failure of NT closure at any level of the body axis leads to neural tube defects (NTDs), in which the neural plate remains open and subsequently degenerates, resulting in loss of neural function below that body level (Copp et al., 2015). Such defects are prominent causes of perinatal mortality and postnatal disability in humans, with NTDs affecting 1 per 1000 pregnancies on average worldwide, and with much higher frequencies in some geographical locations (Zaganjor et al., 2016). In the mouse embryo, NT closure initiates (termed Closure 1) at the hindbrain/cervical boundary at the 6-7 somite stage, with the open neural groove closing *de novo* at the level of the 3^rd^ somite (Sakai, 1989). Initiation of NT closure occurs at a similar stage and somite level in human embryos (O’Rahilly and Müller, 2002). Failure of Closure 1 generates the most severe type of NTD, craniorachischisis (CRN), in which the neural tube remains open from midbrain to low spine (Copp et al., 1994), a defect that comprises around 10% of human NTDs, with a larger proportion in areas of high overall NTD prevalence (Moore et al., 1997).

Initiation of NT closure requires signalling via the planar cell polarity (PCP) pathway, a non-canonical Wnt-dishevelled cascade, that regulates convergent extension (CE). In this process, cells intercalate in the plane of the neural plate or other epithelium, driving medio-lateral narrowing and rostro-caudal elongation, which serves to shape the embryo during and following gastrulation (Skoglund and Keller, 2010). NT closure is disrupted in mice homozygous for mutations in the PCP pathway making this the primary signalling cascade known to be required for initiation of NT closure (Wallingford, 2012). *Vangl2* is the best understood core PCP gene, and homozygosity for the dominant negative *Lp* allele (Strong and Hollander, 1949), or for a null allele (Yin et al., 2012; Ramsbottom et al., 2014), produces CRN. In contrast, single heterozygotes are usually normal except for a looped tail, and occasional spina bifida. Double heterozygosity of *Lp* with other PCP gene mutations generally produces CRN (Murdoch et al., 2014), whereas interaction with genes outside the PCP pathway generates a range of NTDs, including exencephaly and spina bifida. For example, *Lp*/*Grhl3* doubly heterozygous mutants develop severe spina bifida (Stiefel et al., 2007; Caddy et al., 2010; De Castro et al., 2018), while *Lp/Cobl* mutants exhibit exencephaly (Carroll et al., 2003). Hence, the *Lp* mutation is a proven ‘sensitised detector’ that can identify gene-gene, and potentially gene-environmental, interactions that lead to NTDs.

Glycosaminoglycans (GAGs) are linear, unbranched polyanionic molecules of repeating disaccharides that are covalently linked to core proteins to form proteoglycans. GAGs bind and regulate the activity of many secreted factors during development and these interactions depend on the degree of sulfation, epimerisation and acetylation of the chains (Gorsi and Stringer, 2007; Iozzo and Schaefer, 2015). The expression of GAGs is dynamically regulated throughout development (Caterson, 2012), and several studies have shown that the main sulfated GAGs synthesised during primary neurulation in the mammalian embryo are heparan sulfate (HS) and chondroitin sulfate (CS), while the main non-sulfated GAG is hyaluronan (Solursh and Morriss, 1977; Copp and Bernfield, 1988; Yip et al., 2002). These studies suggest a potential role for GAGs in regulating neural tube closure, with support from the finding that the *Lp* mutation interacts genetically with a null mutation in a HS proteoglycan, Syndecan 4 (*Sdc4*), to disrupt spinal neurulation (Escobedo et al., 2013).

Previously, spinal NT closure was found to be affected when wild-type embryos were treated in culture with chlorate, a competitive inhibitor of GAG sulfation (Yip et al., 2002). Moreover, a preliminary study suggested that Closure 1 is also sensitive to chlorate, specifically in *Lp/+* embryos (Escobedo et al., 2013). The current study was therefore designed to investigate in detail the interaction between Vangl2 and GAG chains during initiation of NT closure, and to determine the underlying developmental mechanisms. We find that inhibition of GAG sulfation, or enzymatic removal of sulfated GAG chains, interacts with partial loss of PCP signalling to disrupt Closure 1. Strikingly, however, this effect is not mediated by faulty neuroepithelial convergent extension, which has been found to underlie other instances of CRN in the mouse embryo (Greene et al., 1998; Ybot-Gonzalez et al., 2007; Williams et al., 2014). Hence, additional developmental mechanisms must also be critical for initiation of neural tube closure, and these may be implicated, alongside convergent extension, in the pathogenesis of severe human NTDs.

## RESULTS

The morphology of Closure 1, as visualised in transverse sections, differs from closure at other body levels (Fig. 1A). During cranial and spinal neurulation, the closing neural plate bends focally at either the midline (median hinge point, MHP; Fig. 1A) or at dorsolateral hinge points (DLHPs), or both (Morriss-Kay, 1981; Shum and Copp, 1996). In contrast, the Closure 1 site does not show focal bending at MHP or DLHPs; instead, the neural plate displays a ‘horseshoe’ morphology in cross section (Fig. 1A, B-i,ii; C-i,ii; E-i,ii), in which all parts of the neural plate appear to bend. A further difference is that the closing neural folds are directly flanked by epithelial somites (Fig. 1A, E), not unsegmented cranial or presomitic mesoderm as at other body levels. Hence, the unique morphology and neural plate-mesoderm relationship of the closure initiation site suggests that developmental mechanisms may differ between Closure 1 and other cranial and spinal levels.

**Figure 1.**
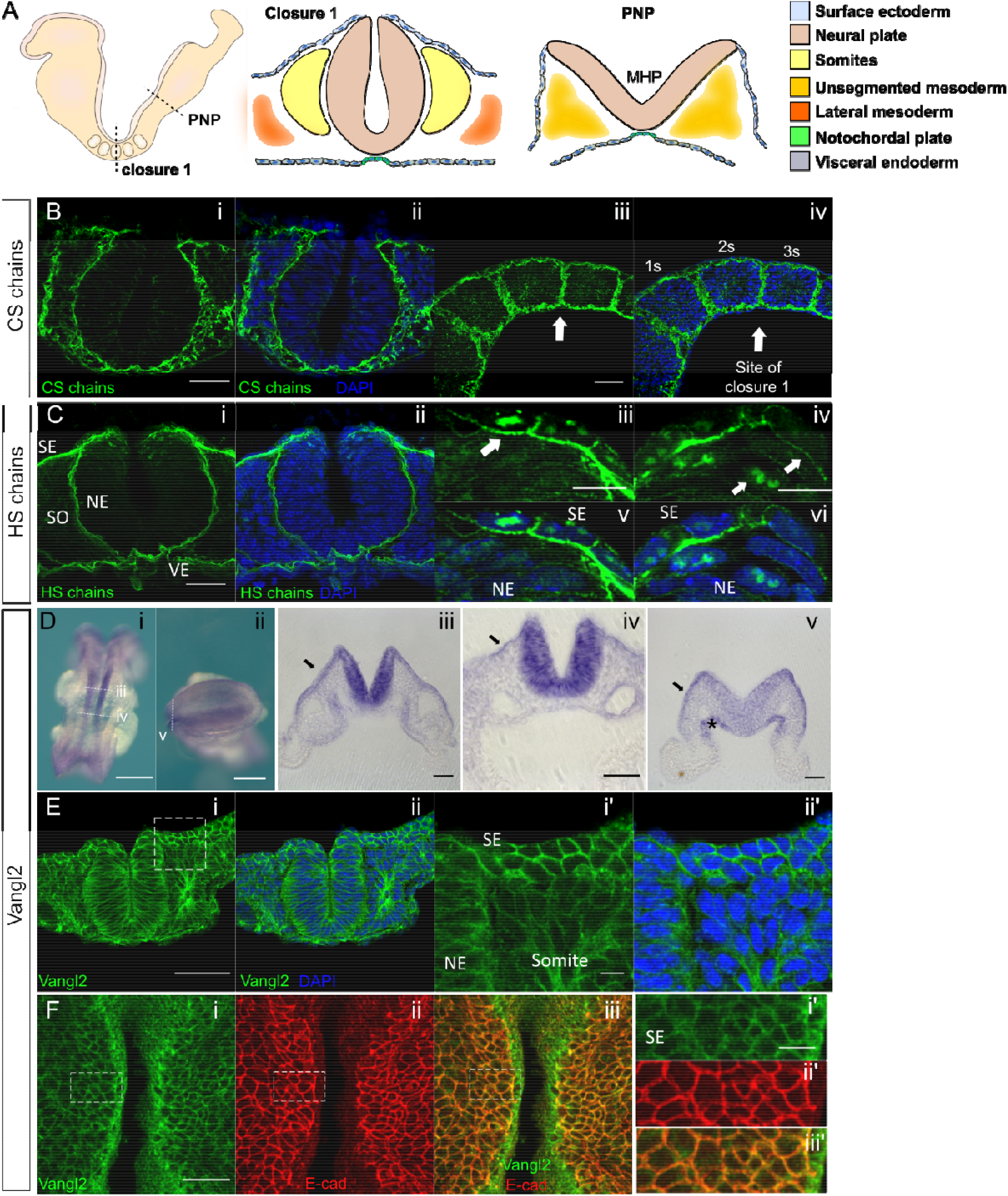
Expression of chondroitin sulfate (CS), heparan sulfate (HS) and Vangl2 at the Closure 1 site of wild-type embryos. (**A**) Schematic representation of an E8.5 mouse embryo (left) showing transverse sections through the Closure 1 region (middle) and posterior neuropore (PNP; right). A median hinge point (MHP) is prominent in the PNP but not at the Closure 1 site. (**B,C**) Immuno-fluorescence staining of CS (B) and HS (C) in transverse (B-i, ii; C-i, ii) and parasagittal (B-iii, iv; C-iii, iv) sections at the Closure 1 site. Both GAG chains localise to the basement membrane of NE and SE (B-i, ii; C-i, ii) (6 somite stage). CS chains are expressed at somite borders (B-iii, iv), with strong staining of HS chains at lateral junctions of SE cells (arrow in C-iii) and weak staining in NE cell membranes (arrows in C-iv). HS chains show nuclear localisation in the dorsal NE and SE at the site of fusion (arrows in C-v, vi). (**D**) *Vangl2* mRNA expression visualised by whole mount *in situ* hybridization (D-i, jj) with transverse vibratome sections (D-iii, iv, v). Transcripts are present throughout the NE, especially at the site of Closure 1 (D-i, iii, iv), and less intensely at the level of the future posterior neuropore (D-ii, v) (6 somite stage). D-ii shows dorsal view of open PNP, anterior to left. *Vangl2* mRNA is also detected in SE (arrows in D-iii, iv, v) and less intensely in visceral endoderm (asterisk in D-v) and paraxial mesoderm, at all levels shown. (**E**) Vangl2 protein is broadly expressed in NE, SE, somitic mesoderm and endoderm (6 somite stage). (**F**) Dorsal view of whole mount embryo (5 somite stage) double immunostained for Vangl2 and E-cadherin (F-i to iii), confirming the presence of Vangl2 in SE (F-i’ to iii’). The SE and apical surface of neural plate was ‘isolated’ virtually using in-house macros. Images acquired by laser-scanning confocal microscopy using oil immersion, deconvoluted post-acquisition, and processed by single z-plane in C-iii to vi and F-i’ to iii’. NE; neuroepithelium; SE: surface ectoderm; SO: somite; VE: visceral endoderm. Scale bars: B, C-i,ii, D-iii-v, E-i,ii = 50 µm; C-iii-vi, E-i’,ii’ = 10 µm; D-i,ii = 250 µm; F-i,ii = 100 µm; F-i’-iii’ = 20 µm.

### Expression of CS and HS chains, and Vangl2 during initiation of neural tube closure

Immunofluorescence analysis on sections of wild-type embryos reveals the distribution of CS and HS chains at the stage and location of Closure 1. Both CS and HS are detected in basement membranes underlying the neuroepithelium (NE), surface ectoderm (SE) and visceral endoderm (Fig. 1B,C). Sagittal sections through the Closure 1 site show strong staining of CS chains at the somite borders and weak staining within the somitic mesoderm (Fig. 1B-iii, iv). HS staining is observed at the basolateral junctions of SE cells, and less intense staining is present around NE cells (Fig. 1C-iii, iv). Nuclear HS staining is observed in the most dorsal NE and SE cells and in some mesodermal cells (Fig. 1C-v, vi). Double immunofluorescence analysis confirmed that both CS and HS chains co-localise with laminin in the SE and NE basement membranes (Fig. S1A,B).

Strong expression of *Vangl2* mRNA is observed at the Closure 1 site (Fig. 1D) (Doudney et al., 2005; Ybot-Gonzalez et al., 2007), with particularly intense signal in NE and at lower intensity expression in somitic, pre-somitic and lateral plate mesoderm. Transcripts are also detected in SE and visceral endoderm (Fig. 1D-iii-v). Immunohistochemistry confirms the widespread expression of Vangl2 protein in cell membranes of all embryonic tissues at the Closure 1 site (Fig. 1E). Double whole mount immunofluorescence with anti-Vangl2 and anti-E-cadherin, an SE marker, verified the presence of Vangl2 in SE during neural tube closure (Fig. S2A), and confirmed the absence of Vangl2 membrane staining in *Vangl2*^*Lp/Lp*^ mutant embryos (hereafter: *Lp/Lp*) (Fig. S2B). The expression pattern of both CS and HS chains is similar between *+/+* and *Lp/Lp* embryos at closure 1 site (Fig. 1B,C vs Fig. S1C,D). Taken together, this analysis reveals that CS and HS chains are co-expressed with Vangl2 in the NE, somitic mesoderm and SE of the Closure 1 region.

### Inhibition of GAG sulfation disrupts Closure 1 in *Vangl2*^*Lp/+*^ and *Vangl2*^*flox/-*^ embryos

Chlorate is a competitive, reversible inhibitor of GAG chain sulfation, commonly used to investigate the role of sulfate groups in proteoglycan function (Conrad, 2001). It has been added to cell and embryo cultures at concentrations up to 30 mM, without apparent adverse effects on GAG or protein synthesis, or cell viability (Greve et al., 1988; Humphries and Silbert, 1988; Yip et al., 2002). We performed a dose-response study in whole embryo culture, to identify a minimum concentration of chlorate that would affect Closure 1 in *Vangl2*^*+/+*^ and *Vangl2*^*Lp/+*^ (hereafter: *+/+* and *Lp/+* respectively) embryos without toxic effects on embryonic growth and development (Table S1). Closure 1 was significantly inhibited (entirely open NT) in the majority of *Lp/+* embryos treated with 10 and 20 mM chlorate, whereas most wild-type littermates underwent successful Closure 1 at the same inhibitor concentrations (Fig. 2). The effect of chlorate on Closure 1 showed a similar dose-response in *Lp/+* embryos after a single generation outcross to BALB/c, as well as on the original CBA/Ca genetic background (Fig. 2A,B). Chlorate at 10 mM had no adverse effects on embryo health parameters and was used in subsequent culture experiments with *+/+* and *Lp/+* embryos on the CBA/Ca genetic background.

**Figure 2.**
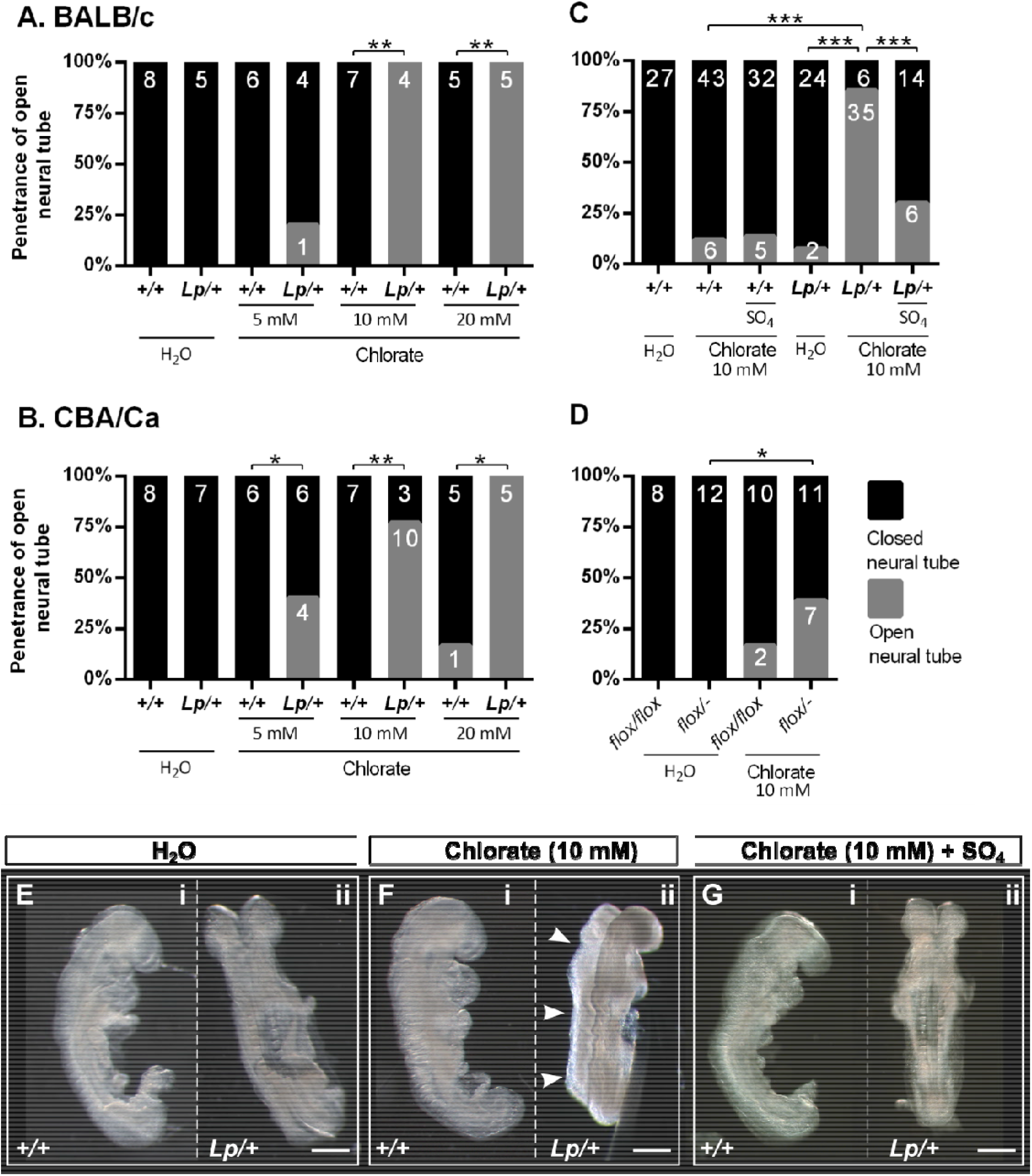
Chlorate induces failure of Closure 1 in *Vangl2*^*Lp/+*^ and *Vangl2*^*flox/-*^ embryos. (**A,B**) Dose-response study of chlorate effect on *+/+* and *Lp/+* embryos in embryo culture. Experimental litters were generated by a single outcross of *Vangl2*^*Lp/+*^ males to BALB/c females (A) or by *Vangl2*^*Lp/+*^ x *Vangl2*^*+/+*^ matings within the CBA/Ca genetic background (B). Percentage of open NT (grey bar sectors) at the Closure 1 site was determined in cultures containing 5, 10 or 20 mM chlorate, or water as control. Embryo numbers are shown on bars. Treatment with chlorate at 5 mM (CBA/Ca only) and at 10 and 20 mM (both BALB/c and CBA/Ca) prevents Closure 1 in a significantly greater proportion of *Lp/+* than *+/+* embryos. Closure 1 is successful in 100% of water-treated controls, both *+/+* and *Lp/+* genotypes. (**C**) Rescue experiment (CBA/Ca background) in which treatment with sodium sulfate (SO_4_) significantly reduces the Closure 1 failure seen in *Lp/+* embryos treated with 10 mM chlorate. (**D**) Chlorate significantly inhibits Closure 1 in *Vangl2*^*flox/-*^ embryos, compared with *Vangl2*^*flox/flox*^, although the penetrance of Closure 1 failure after 10 mM chlorate is significantly lower in *Vangl2*^*flox/-Lp/+*^ embryos (7/18) than in *Vangl2* embryos (CBA/Ca: 35/41; Fig 2C; p < 0.001). (**E-G**) Typical embryonic morphology after 24 h culture. *Vangl2*^*+/+*^ embryos show completion of Closure 1 in all treatment groups (E-i, F-i, G-i). *Vangl2*^*Lp/+*^ water-treated (E-ii) and chlorate + sulfate-treated embryos (G-ii) also show Closure 1, whereas a chlorate-treated *Vangl2*^*Lp/+*^ embryo shows entirely open neural tube (arrowheads in F-ii). *p < 0.05; **p < 0.01;***p < 0.001. Scale bars: 0.5 mm in E-G.

A rescue experiment was performed in which embryos received 10 mM sodium sulfate in addition to 10 mM chlorate in the culture medium. Exogenous sulfate is known to restore GAG sulfation in a dose-dependent manner, without competitively inhibiting chlorate (Humphries and Silbert, 1988). The frequency of Closure 1 failure resulting from chlorate exposure in *Lp/+* embryos decreased significantly from 85% in the absence of sodium sulfate to 30% in its presence (Fig. 2C), without any adverse effects on embryo morphology (Fig. 2E-G). Importantly, the percentage of *Lp/+* embryos with open NT did not differ significantly between the sulfate rescue and water control groups (Fig. 2C), demonstrating that exogenous sulfate is able to rescue most *Lp/+* embryos from chlorate-induced Closure 1 failure.

The effect of chlorate was tested independently in a targeted transgenic line *Vangl2*^*flox/-*^. Previously, global loss of Vangl2 using this knockout allele was shown to recapitulate the *Lp/Lp* phenotype (Ramsbottom et al., 2014). Both *Vangl2*^*flox/flox*^ and *Vangl2*^*flox/-*^ embryos cultured under control conditions completed Closure 1 (Fig. 2D). In contrast, a proportion of *Vangl2*^*flox/-*^ embryos failed in Closure 1 when treated with 10 mM chlorate, whereas *Vangl2*^*flox/flox*^ embryos were not significantly affected by the inhibitor (Fig. 2D). The penetrance of Closure 1 defect in chlorate-treated *Vangl2*^*flox/-*^ embryos (39%; 7/18) was significantly lower than in chlorate-treated *Lp/+* embryos on the CBA/Ca background (Fig 2C: 77%; 35/41; p < 0.001), consistent with previous findings that *Lp* has more severe (likely dominant negative) effects on PCP signalling compared with *Vangl2*-null alleles (Yin et al., 2012).

We asked whether a developmental ‘window’ exists when Closure 1 is susceptible to chlorate treatment. Somite number reached at the end of culture did not differ between genotype or treatment groups (Fig. S3A). Somite stage at the start of culture was calculated by subtracting the expected number of new somites formed during the culture period (based on formation of one somite every 2 h) from the number of somites recorded at the end of culture (Fig. S3B). The great majority of *+/+* embryos completed NT closure, even when treated with chlorate. In contrast, most *Lp/+* embryos failed in NT closure after exposure to 10 mM chlorate, and this comprised 92% (22/24) of those treated from the 0-1 somite stage, 83% (10/12) from the 2-3 somite stage and 60% (3/5) from the 4-5 somite stage (p > 0.05; Fig. S3C). Hence, chlorate can inhibit Closure 1 even if administered only a few hours before the event.

Immunofluorescence analysis was performed to determine the effect of chlorate treatment on the distribution of sulfated GAGs in the cultured embryos. Wild-type and *Lp/+* embryos cultured under control conditions exhibit strong CS and HS staining (Fig. 3A,D,G,J), similar to the immunostaining of wild-type embryos prior to Closure 1 (Fig. 1A,B). In contrast, CS and HS are dramatically reduced in chlorate-treated *+/+* and *Lp/+* embryos (Fig. 3B,E,H,K). Addition of sulfate rescues the expression pattern of CS and HS chains in chlorate-treated *+/+* and *Lp/+* embryos (Fig. 3C,F,I,L), generating a staining pattern closely resembling the controls (Fig. 3A,D,G,J). This chlorate-specific reduction in GAG sulfation correlates closely with Closure 1 inhibition in *Lp/+* embryos, although notably the inhibition of CS and HS immunostaining is present also in +/+ embryos, which are not adversely affected by chlorate in NT closure.

**Figure 3.**
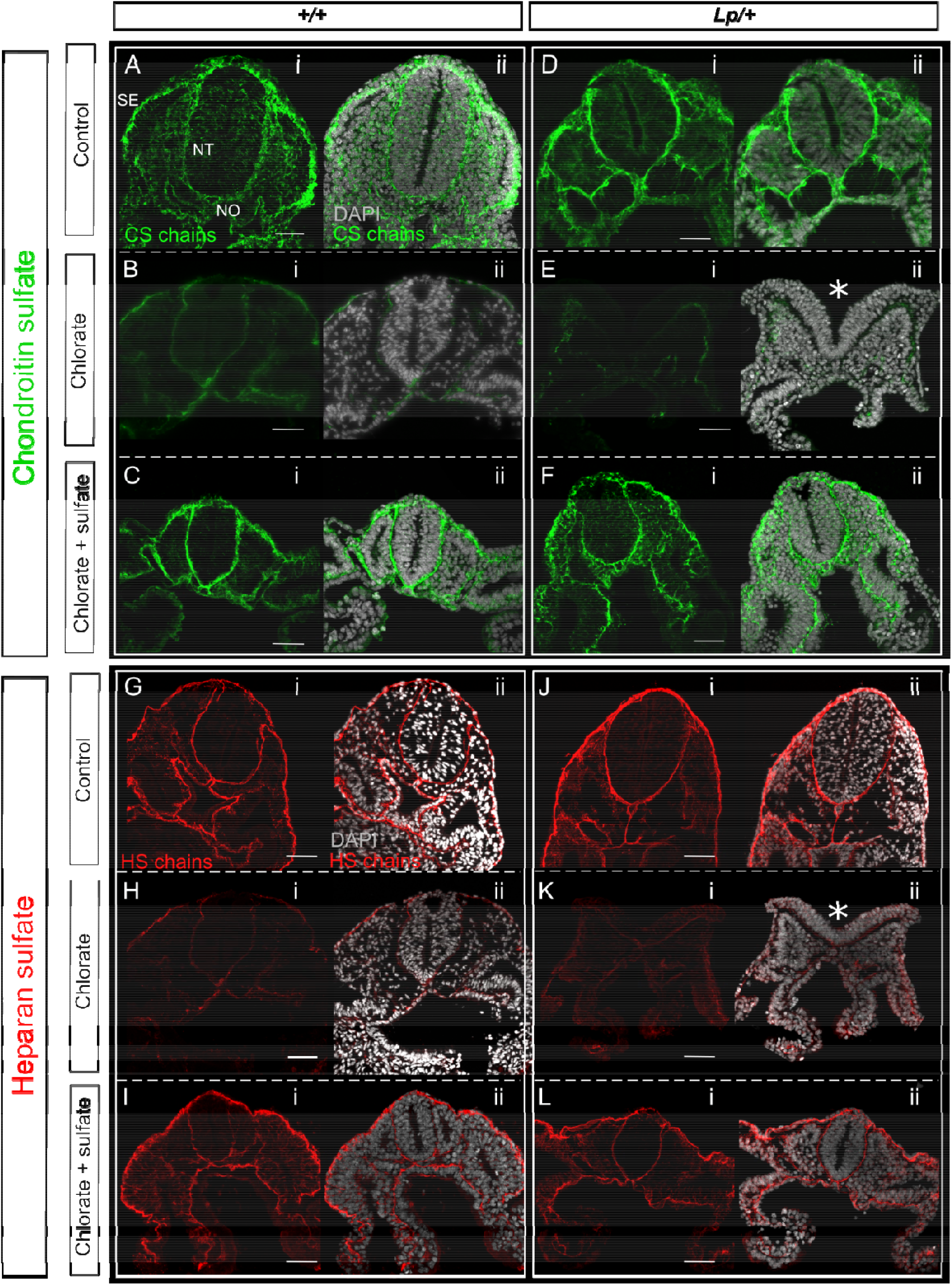
Chlorate reduces sulfation of CS and HS chains in cultured embryos. Immunofluorescence analysis of CS (A-F) and HS (G-L) chains in embryos cultured in the presence of water (control), 10 mM chlorate, or 10 mM chlorate plus sodium sulfate. (**A,D,G,J**) Sections from control *+/+* and *Lp/+* embryos have normal distribution of CS and HS chains, which are localised to the BM underlying SE and surrounding the NT and notochord (NO); CS staining is also detected around mesenchymal cells. (**B,E,H,K**) Chlorate treatment dramatically reduces the staining of CS and HS chains in both genotypes and leads to failure of Closure 1 in *Lp/+* embryos (asterisks in E-ii, K-ii: open NT). (**C,F,I,L**) Staining of CS and HS chains in *+/+* and *Lp/+* embryos from the rescue group (chlorate plus sulfate) appears similar to the control group. Note the closed NT in *Lp/+* embryos. Stages shown: 12-14 somites. Scale bar: 50 µm.

### Enzymatic cleavage of GAG chains recapitulates the phenotype of chlorate treatment in *Vangl2*^*Lp/+*^ embryos

Chlorate affects the sulfation of both HS and CS chains (Yip et al., 2002). To address whether one or other type of GAG chain is particularly required for the closure initiation, *+/+* and *Lp/+* embryos with 0-5 somites were cultured following intra-amniotic injection of specific GAG-degrading enzymes: chondroitinase ABC (Chr.ABC) for CS, or heparitinase III (Hep.III) for HS. Specificity of these enzymes was demonstrated on embryo sections immunostained for CS or HS, following enzyme treatment in embryo culture (Fig. S4). All *+/+* and *Lp/+* embryos in buffer-treated cultures completed Closure 1, and exhibited normal morphology (Fig. 4A,B), whereas NT closure was significantly affected in *Lp/+* embryos treated with Hep.III (7/7) or Chr.ABC (7/10), with Closure 1 failure and entirely open NTD in both cases (Fig. 4A, C, D). A small number of *+/+* embryos failed in Closure 1 after treatment with either Hep.III or Chr.ABC, although the frequency of NT closure failure was not significantly different from +/+ buffer-treated controls.

**Figure 4.**
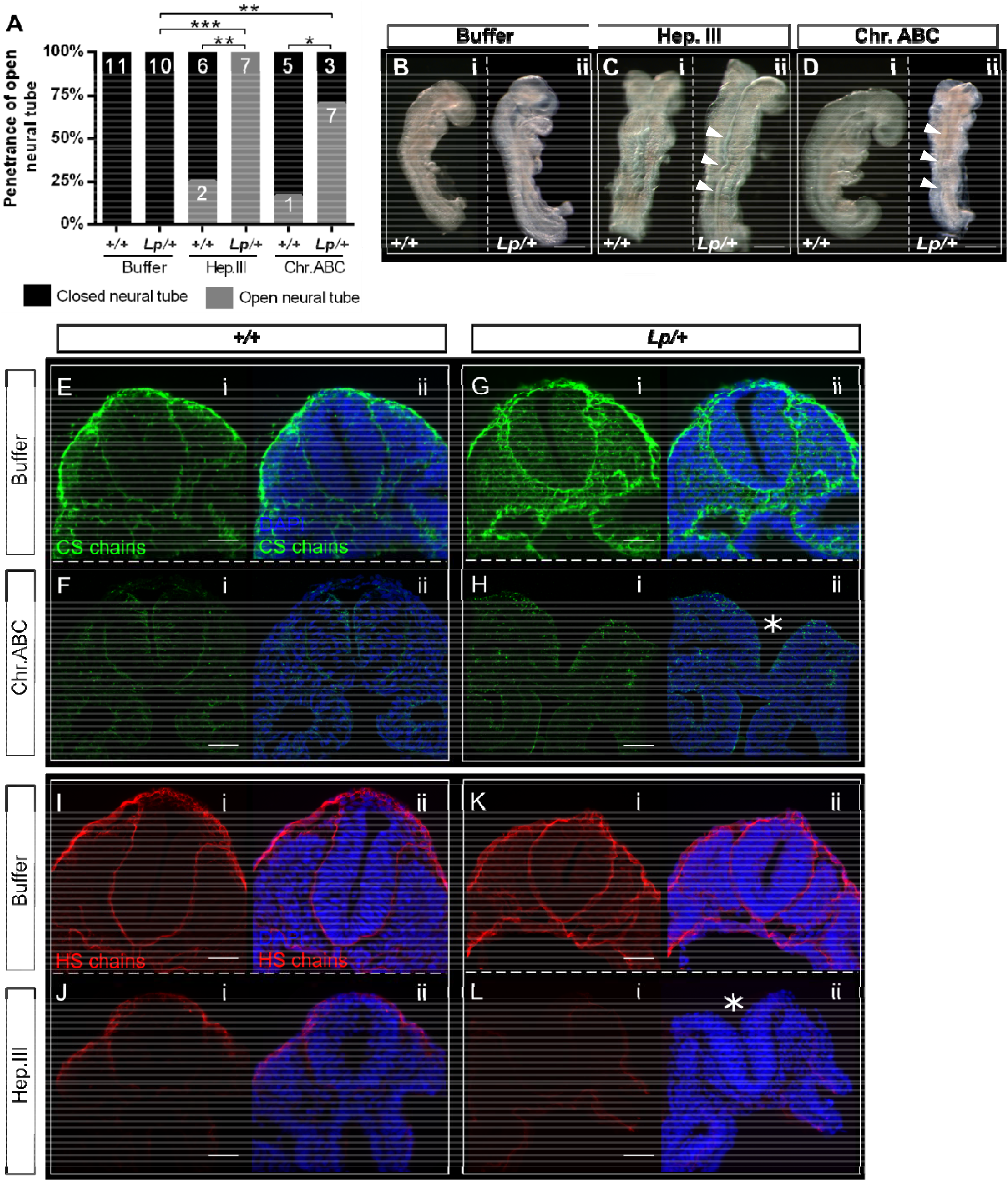
Enzymatic cleavage of CS and HS chains induces Closure 1 failure in *Lp/+* embryo cultures. Heparitinase III (Hep.III, to cleave HS chains, 5U/ml) or chondroitinase ABC (Chr.ABC, to cleave CS chains, 2U/ml) were injected into the amniotic cavity of E8.5 embryos at 0-5 somite stage (prior to Closure 1) at time 0, and injection was repeated after 4 h of culture. Embryos were cultured for 24 h in total. (**A**) All *+/+* and *Lp/+* embryos from the control group (enzyme buffer only) completed Closure 1. Both Hep.III and Chr.ABC significantly inhibited Closure 1, producing open NT (grey bar sectors) in *Lp/+* embryos whereas *+/+* embryos remained largely unaffected. Embryo numbers are shown on bars. (**B-D**) Examples of embryos after culture, showing *+/+* and *Lp/+* buffer-treated embryos with closed NT (B-i, ii), *+/+* embryos with closed NT despite treatment with enzymes (C-i, D-i), and *Lp/+* embryos with failed Closure 1 after Hep.III and Chr.ABC (arrowheads in C-ii and D-ii). (**E,G,I,K**) Both *+/+* and *Lp/+* embryos from the control group (buffer) show the expected distribution of CS/HS chains after culture (compare with Fig. 3A,D,G,J). (**F,H**) Chr.ABC injection dramatically reduces the staining of the CS chains in both genotypes. (**J,L**) *+/+* and *Lp/+* embryos injected with Hep.III display very weak staining of HS chains. Chr.ABC does not affect HS staining, nor does Hep.III affect CS staining (data not shown). Somite stages: B-i, 12; B-ii, 13; C-i, 14; C-ii, 15; D-i, 14; D-ii, 10; E-G, 12; H, 11; I, 13; J-L, 12. *p < 0.05; **p < 0.01;***p < 0.001. Scale bars: 0.5 mm in B-D, 50 µm in E-L.

To confirm the effect of enzymatic treatment on GAG chains, immunofluorescence analysis was carried out on the cultured embryos. Both +/+ and *Lp*/+ embryos in control cultures show strong immunostaining staining of CS and HS chains (Fig. 4E,G,I,K). In contrast, the staining of CS chains was almost absent in the Chr.ABC treated *+/+* and *Lp/+* embryos (Fig. 4F,H), while exposure to Hep.III dramatically reduced HS staining, with only some SE basement membrane staining persisting in *+/+* embryos (Fig. 4J,L). Hence, enzymatic cleavage of CS and HS chains each recapitulates the Closure 1 phenotype of chlorate-treated *Lp/+* embryos suggesting that both chain types are required for the initiation of NT closure.

### Closure 1 failure in chlorate-treated *Lp/+* embryos is not due to faulty convergent extension

Mouse embryos homozygous for PCP mutations exhibit abnormal convergent extension (CE) cell movements, preceding failure of Closure 1 (Greene et al., 1998; Ybot-Gonzalez et al., 2007; Williams et al., 2014). Moreover, CRN, the severe NTD that results from Closure 1 failure, has been found almost exclusively in association with a short, wide body axis, indicative of defective CE (Curtin et al., 2003; Wang et al., 2006; Yen et al., 2009; Murdoch et al., 2014). We hypothesised that the Closure 1 defects observed in *Lp/+* embryos after chlorate treatment result from enhancement of a heterozygous predisposition to faulty CE. Vital labelling of the embryonic midline at the late gastrulation stage has shown that *Lp/Lp* mutants fail in CE both in NE and axial mesoderm, prior to the stage of Closure 1 (Ybot-Gonzalez et al., 2007). Here, we used the same labelling technique to investigate the effect of chlorate on CE in *Lp/+* embryos. *Lp/Lp* mutants served as ‘positive controls’, as they were expected to show reduced CE even under normal culture conditions.

The embryonic midline was labelled by focal injection of DiO into the node at E8.5 in *+/+, Lp/+* and *Lp/Lp* embryos, prior to culture with or without chlorate (Fig. 5A). Analysis of transverse embryo sections immediately after labelling (time 0) revealed that both node and floor plate were successfully labelled with DiO in all three genotypes (Fig. 5A-i’-iii’). Analysis after 24 h culture revealed that both *+/+* and *Lp/+* embryos from the water-treated control group exhibit rostrally-directed midline extension of DiO-labelled cells (Fig. 5B,C). Moreover, chlorate-treated *+/+* and *Lp/+* embryos also exhibit rostrally-directed midline extension of DiO-labelled cells (Fig. 5E,F), closely resembling the water-treated controls. In contrast, *Lp/Lp* embryos display very limited midline extension, whether treated with water (Fig. 5D) or chlorate, with the latter having no discernible effect on midline extension in *Lp/Lp* embryos (Fig. 5G). Labelled cells were persistently located at the site of node injection in *Lp/Lp* embryos, as described previously (Ybot-Gonzalez et al., 2007). Transverse sections demonstrate that, following 24 h culture, labelled cells can be detected extending along the midline, from the caudal region to upper trunk in *+/+* and *Lp/+* embryos, with no obvious difference between water- or chlorate-treatments (Fig. 5B-ii, C-ii, E-ii, F-ii). In contrast, midline extension of DiO labelled cells was not detected in the notochord and floor plate of *Lp/Lp* embryos, either water- or chlorate-treated (Fig. 5D-ii and G-ii).

**Figure 5.**
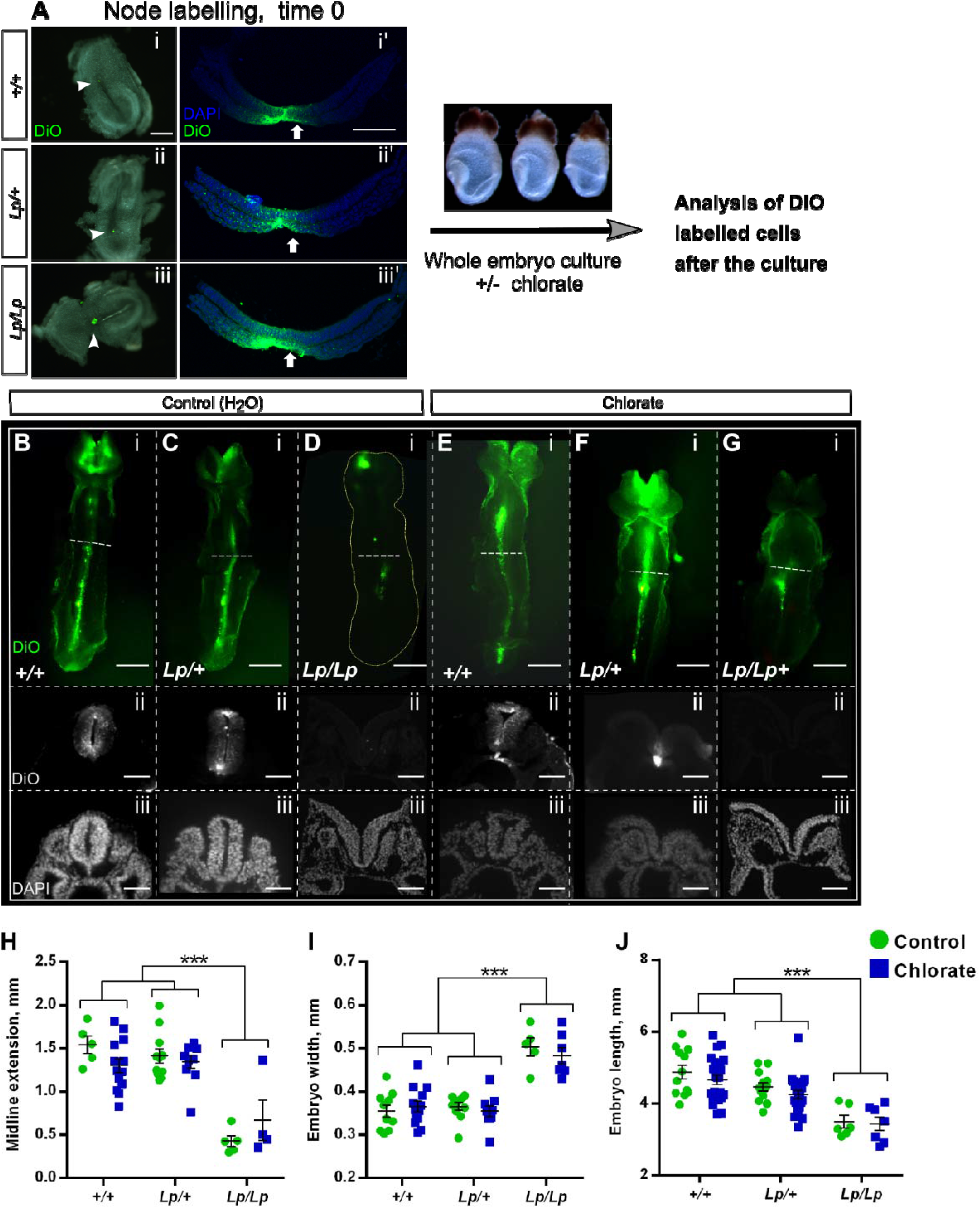
Chlorate prevents Closure 1 without affecting midline extension of *Lp/+* embryos. (**A**) The node of E8.5 embryos from *Vangl2*^*Lp/+*^ x *Vangl2*^*Lp/+*^ matings (0-5 somite stage, blind to genotype) was labelled by microinjection of the lipophilic fluorescent dye (DiO) (A-i-iii). Embryos were then randomly allocated to 10 mM chlorate or water treatment and cultured for 24 h. Transverse sections through embryos at time 0 show the node and floor plate are successfully labelled with DiO in all three genotypes (A-i’-iii’). (**B-G**) Ventral view (rostral to the top) and transverse sections (trunk region) of DiO injected embryos, after 24 h culture. Control (water) and chlorate-treated *+/+* and *Lp/+* embryos (B, C, E, F) exhibit marked midline extension of DiO labelled cells, as detected in both whole mount and sections. Sections reveal failed Closure 1 in chlorate-treated *Lp/+* embryos (F-ii, iii). In contrast, *Lp/Lp* embryos from both treatment groups display very limited midline extension of DiO labelled cells (D-i; G-i), and fail in Closure 1 (D-ii, iii; G-ii, iii). Note: DiO labelling of the cranial region is non-specific, due to release of DiO into the amniotic cavity during labelling. (**H**) Midline extension measurements: points represent distance DiO labelled cells have extended along the caudal-to-rostral axis in individual embryos, with mean +/- SEM also shown. *Lp/Lp* embryos exhibit significantly less midline extension than other genotypes. Chlorate-treated *Lp/+* embryos do not differ from *+/+* (water or chlorate) or *Lp/+* (water) groups. (**I,J**) Embryo width (I) and length (J) measurements reveal a significantly wider and shorter body axis in *Lp/Lp* embryos than in other genotypes, irrespective of water/chlorate treatment. Width and length of *Lp/+* embryos does not differ significantly between water and chlorate groups, nor do these values differ from *+/+* embryo measurements. Data points are individual measurements, with mean +/- SEM values also shown. *** p < 0.001. Scale bars: A-i-iii = 200 µm; A-i’-iii’ = 100 µm; B-i – G-i = 0.5 mm; B-ii, iii – G-ii, iii = 50 µm.

Midline extension was quantified by measuring from the caudal end of the embryo to the rostral limit of continuous midline DiO signal, which typically ended around heart level (Fig. 5H). Chlorate treatment did not affect midline extension overall (ANOVA, p = 0.92), and there was no statistically significant interaction between genotype and treatment (p = 0.21). Midline extension varied significantly between genotypes (p < 0.001), as a result of the greatly reduced extension in *Lp/Lp* embryos, whereas *+/+* and *Lp/+* genotypes did not differ significantly from each other (p = 0.41).

We evaluated embryo width and length after culture, as a further measure of CE (Fig. 5I,J). Chlorate treatment did not significantly affect embryo width and length overall (p = 0.28 and p = 0.44 respectively) and there was no statistically significant interaction between genotype and treatment (p = 0.27 for width; p = 0.89 for length). Embryo width and length varied significantly between genotypes, with both *+/+* and *Lp/+* embryos exhibiting smaller width and increased length compared with *Lp/Lp* littermates (p < 0.001 for both). Whether treated with chlorate or water, *+/+* and *Lp/+* embryos do not differ from each other for either width or length (p > 0.05). In conclusion, chlorate treatment does not appear to disrupt the process of neuroepithelial CE in *Lp/+* embryos, even though these fail in Closure 1 and develop CRN at high frequency. As expected, *Lp/Lp* embryos exhibit defective CE, whether treated with water or chlorate.

### Effect of chlorate treatment on neural plate morphology in the Closure 1 region

To further investigate the embryonic mechanisms leading to failure of NT closure in chlorate-treated *Lp/+* embryos, we examined the morphology of mid-axial tissues during the onset of Closure 1. Cultures were started at the 0-4 somite stage (prior to Closure 1) and continued for 8 h (as opposed to 24 h cultures in previous experiments), in the presence of 10 mM chlorate or water as control. Neural plate morphology of the Closure 1 region was then examined in *+/+* and *Lp/+* embryos at the 5, 6, 7 and 8 somite stage. Embryos were whole-mount stained with CellMask(tm) to label all tissues, followed by confocal imaging and ‘re-slicing’ in Fiji to obtain virtual transverse sections of the Closure 1 region, at the level of the third somite (Fig. 6).

**Figure 6.**
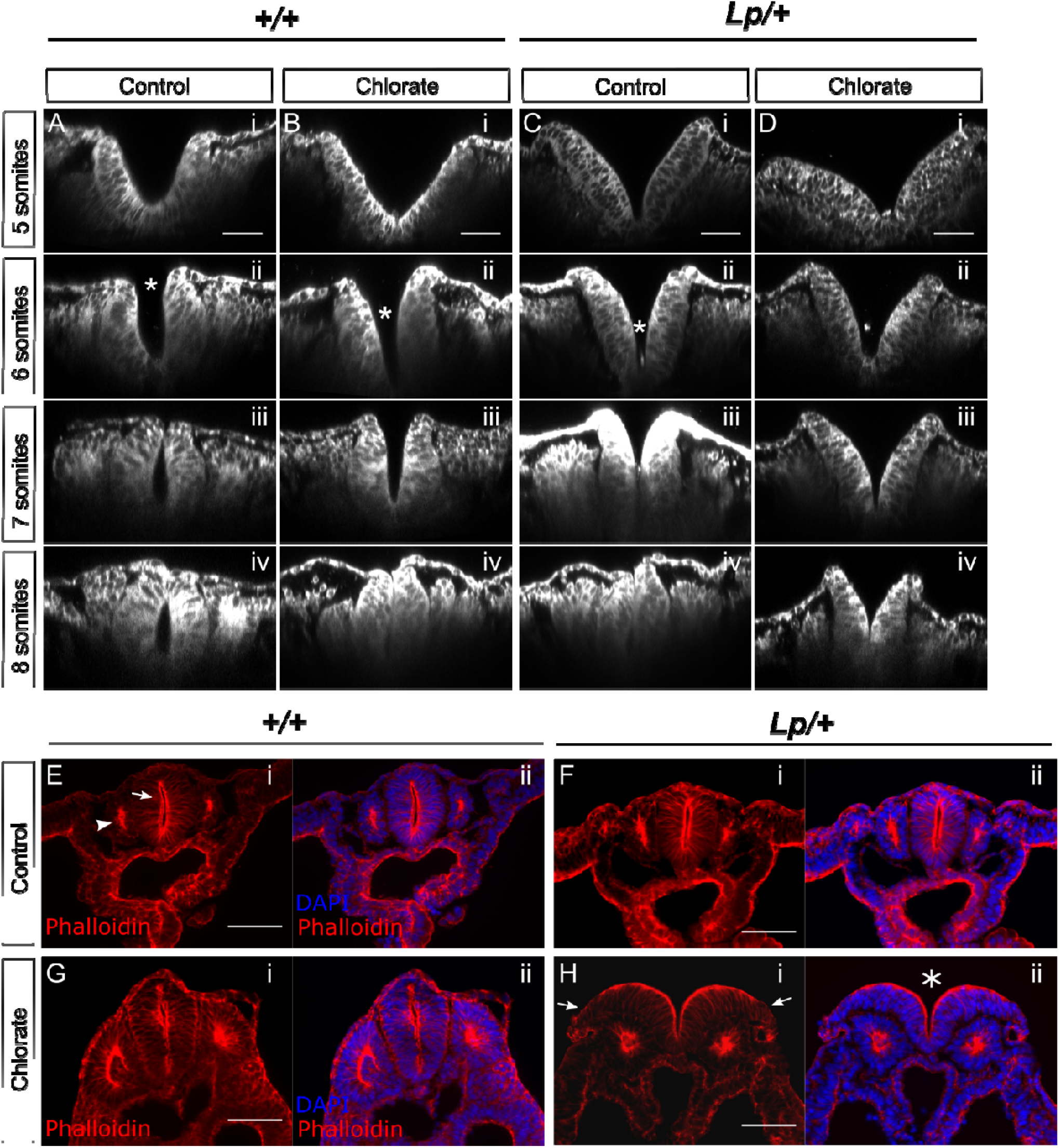
Chlorate alters neural plate morphology but not overall F-actin distribution in cultured embryos. (**A-D**) Embryos were cultured for 8 h with addition of 10 mM chlorate or water as control, fixed, stained with CellMask™ and imaged using confocal microscopy for morphological analysis. Images were re-sliced in Fiji to obtain transverse sections of the Closure 1 region, at the level of the third somite. *+/+* (A,B) and *Lp/+* (C,D) embryos with 5, 6, 7 or 8 somites are shown (i to iv for each genotype/treatment combination). Closure 1 is normally completed from the 6 somite stage onwards. In control (water-treated) *+/+* embryos, the neural plate adopts an increasingly horseshoe shape, concave inwards, with fusion evident dorsally from 7 somites (A). Chlorate delays this transition in *+/+* embryos, with an initially V-shaped morphology, but closure is achieved by 8 somites when the neural tube appears largely normal (B). Water-treated *Lp/+* embryos resemble chlorate-treated *+/+* embryos, and achieve closure by 8 somites (C). Chlorate-treated *Lp/+* embryos exhibit a persistently V-shaped neural plate with convex curvature, in which the dorsal aspects of the neural folds fail to converge and fusion fails (D). Asterisks in A-D-ii indicate sites of initial contact between neural folds. (**E-H**) Phalloidin staining to detect F-actin distribution in transverse sections of the Closure 1 region of embryos cultured for 24 h. F-actin is enriched at the apical surface of NE (arrow in E-i) and apically within the epithelial somites (arrowhead in E-i). Although failure of NT closure is seen in chlorate-treated *Lp/+* embryos (asterisk in H-ii), the only obvious difference from *+/+* (E, G) and water-treated *Lp/+* embryos (F) is reduced intensity of phalloidin staining at the lateral edges of the open neural folds (arrows in H-i)(n = 4 embryos each). Scale bar: A-D = 50 µm; E-H = 100 µm.

Confocal microscopy revealed the typical ‘horseshoe’ morphology of the NE at the Closure 1 site in *+/+* embryos in the control treatment group. All regions of the neural plate appear to bend in embryos with 5-7 somites so that the neural fold tips come together and fuse in the dorsal midline at the 7 somite stage. A completely closed neural tube is present by 8 somites (Fig. 6A). A different morphology was noted in chlorate-treated *+/+* embryos, in which a well-defined bend occurred at the neural plate midline at the 5 and 6 somite stages (Fig. 6B). This resembles the MHP that characterises closure at upper spinal levels, following completion of Closure 1 (Shum and Copp, 1996). Hence, the ‘horseshoe’ morphology was less obvious in these embryos. Nevertheless, their neural folds invariably came into contact and appeared to fuse just below the dorsal tips (asterisk in Fig. 6B-ii), thereby completing Closure 1 by the 8 somite stage.

*Lp/+* embryos in the water-treated group also exhibited a ‘V’-shaped neural plate at the 5 and 6 somite stage (Fig. 6C), with a sharper midline bend than in chlorate-treated *+/+* embryos. Their neural folds elevated and came into contact apparently in a ventral-to-dorsal sequence within the NE (Fig. 6C-ii, iii), rather than by initial contact between the neural fold tips (Fig. 6A-ii, iii). Closure 1 was completed by the 8 somite stage. The neural plate of chlorate-treated *Lp/+* embryos had a similar morphology to water-treated *Lp/+* embryos at the 5 and 6 somite stage (Fig. 6D), but the distance between neural fold tips was greater, as a result of bilateral convexity of the NE. This was observed at all stages up to and including 8 somites, and hence Closure 1 failed in these embryos.

Measurements of the distance between the neural fold tips confirmed that both chlorate-treated *+/+* and water-treated *Lp/+* embryos exhibit delayed Closure 1 compared with normally developing, water-treated *+/+* embryos. Moreover, chlorate-treated *Lp/+* embryos show no reduction in inter-fold distance with increasing somite number, and fail in Closure 1 (Fig. S5).

### Chlorate does not perturb the distribution of F-actin in the neural folds

The finding that chlorate-treated *Lp/+* embryos exhibit convex, rather than concave, neural folds suggested a possible mechanism whereby chlorate may disrupt the NE actin cytoskeleton leading to failure of Closure 1. Apically arranged F-actin microfilaments are well known to participate in neural tube closure, at least in part by biomechanically stabilising the apical NE and promoting concave curvature (Sawyer et al., 2010; Suzuki et al., 2012). To examine this possibility, we performed phalloidin staining to reveal F-actin distribution in the Closure 1 region of embryos cultured for 24 h. Transverse sections demonstrate an F-actin enrichment at the apical surface of the NE in *+/+* and *Lp/+* embryos from the water-treated control group (Fig. 6E,F), with prominent staining also of the central (apical) region of the epithelial somites that flank the closed neural tube. Chlorate-treated *+/+* embryos display a similar F-actin distribution on the apical aspects of the NE and epithelial somites (Fig. 6G). In chlorate-treated *Lp/+* embryos, the lateral-most parts of the open neural folds show a reduced intensity of phalloidin staining (arrows in Fig. 6H), but the majority of each neural fold has prominent F-actin localisation apically, including in regions of the greatest convex curvature. We conclude that there is no major alteration in F-actin distribution that can account for the failure of Closure 1 in chlorate-treated *Lp/+* embryos.

### Possible somite-mediated mechanism of Closure 1 failure in chlorate-treated *Lp/+* embryos

Closure 1 is unusual in mammalian neurulation as it occurs at a rostro-caudal level where the neural folds are flanked by epithelial somites. Most of spinal closure, and all of cranial closure, occur when the neural tube is adjacent to unsegmented presomitic or cranial mesoderm, respectively. Hence, we investigated whether the somites might play a functional role in promoting Closure 1, with possible defects resulting from chlorate treatment. Somite length (rostro-caudal) and width (mediolateral) were measured in confocal images at the level of the third and fourth somites of embryos cultured for 8 h, to the 6-7 somite stage, when onset of Closure 1 is underway (Fig. 7A). Statistical analysis revealed that somite length is significantly reduced by chlorate treatment relative to water-treated control embryos (ANOVA; p = 0.023; Fig. 7B). Somite length is also reduced in *Lp/+* relative to *+/+* embryos, a difference that just reaches significance (p = 0.049). The interaction between genotype and treatment was not statistically significant for somite length.

**Figure 7.**
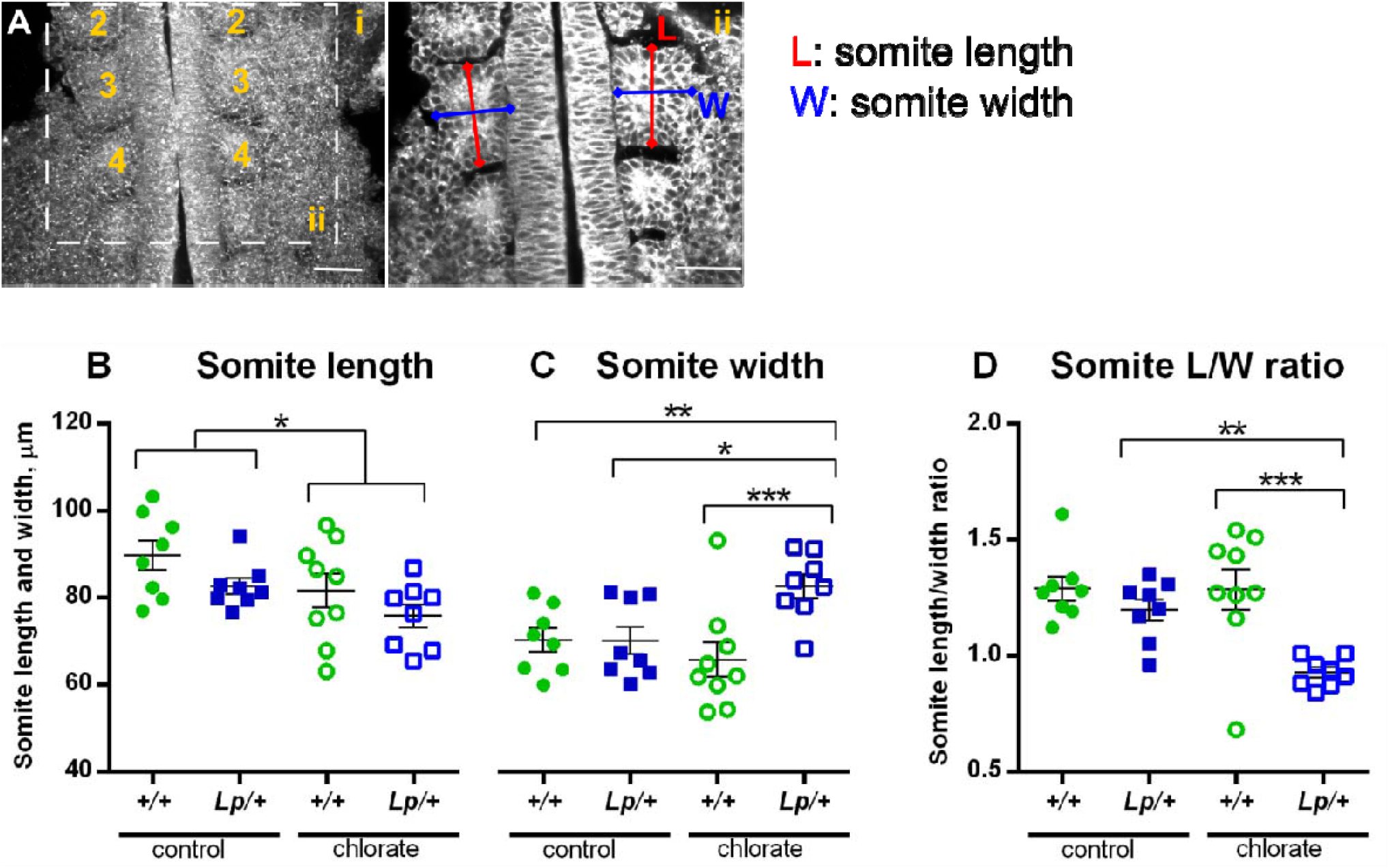
Somite morphology at the Closure 1 site is altered by chlorate treatment. Embryos were cultured for 8 h from the 0-5 somite stage, with addition of 10 mM chlorate or water as control, and prepared for morphological analysis as in Fig. 6A-D. (**A-i**) Z-projection of *+/+* embryo at 7 somite stage from control group. Rostral at top; somites numbered sequentially. Images were re-sliced in Fiji (from dorsal to ventral surface) to obtain horizontal sections through the somite row. (**A-ii**) Single Z-plane through the Closure 1 region. Length (L; rostrocaudal orientation) and width (W; mediolateral orientation) measurements were taken half way though the somite, at the level of 3rd/4th somites. Two somite pairs (4 individual somites) were measured per embryo. (**B-D**) Somite length, width and length/width ratio in control and chlorate-treated *+/+* and *Lp/+* embryos, at the 6-7 somite stage. Individual points on the graphs are measurements averaged over four somites for each embryo, with mean +/- SEM of embryo replicates. Note that somite width is significantly increased, and length/width ratio significantly reduced, in chlorate-treated *Lp/+* embryos. * p < 0.05; ** p < 0.01;*** p < 0.001.

When considering somite width, there is a significant genotype-treatment interaction (p = 0.015) in which chlorate treatment increased somite width of *Lp/+* embryos relative to chlorate-treated *+/+* embryos and also relative to *Lp/+* embryos from the water-treated control group (Fig. 7C). Somite length/width (L/W) ratio showed a particularly strong genotype-treatment interaction (p = 0.034), with chlorate-treated *Lp/+* embryos having a significantly reduced L/W ratio compared with the other genotype/treatment groups (p = 0.017) (Fig. 7D). The L/W ratios of *+/+* embryos and water-treated *Lp/+* embryos are 1.2-1.3, indicating their somites are elongated rostro-caudally, whereas L/W ratio in chlorate-treated *Lp/+* embryos is close to 1.0, showing these embryos to have somites that are as wide as they are long. Hence, an abnormality of somite morphology correlates with propensity to Closure 1 failure in chlorate-treated *Lp/+* embryos.

## DISCUSSION

In the present study, we asked whether interactions between the PCP signalling pathway and molecules of the cell surface and extracellular matrix (ECM) may play a role in the initial event of mouse NT closure. We particularly focus on sulfated GAG chains that decorate proteoglycans. Heterozygosity for the *Vangl2*^*Lp*^ PCP mutant allele is ordinarily compatible with initiation of NT closure, but we find a high frequency of Closure 1 failure when synthesis or presence of sulfated GAG chains is also abolished. Hence, partial loss of PCP function can give rise to a severe NTD when combined with loss of GAG sulfation during early embryogenesis.

In zebrafish and *Xenopus*, the axial tissues including neural plate, notochord and mesoderm undergo PCP-dependent mediolateral convergence and rostro-caudal extension (Tada and Heisenberg, 2012). Moreover, ECM proteins and their receptors play a role in CE, with perturbation of ECM components giving rise to defective CE (Topczewski et al., 2001; Skoglund and Keller, 2010). Indeed, PCP signalling and ECM can integrate at the molecular level to regulate CE during neurulation of lower vertebrates. For example, recruitment of *Dsh* (a key intracellular PCP component) to the cell membrane is dependent on fibronectin/integrin and syndecan-4/fibronectin interactions in *Xenopus* CE (Munoz et al., 2006).

There have so far been few studies of PCP signalling in the context of ECM dysfunction in higher vertebrate CE. Our finding of Closure 1 failure in *Lp/+* embryos with under- or non-sulfated GAGs, led to an initial hypothesis of defective CE, with PCP heterozygosity summating with ECM disturbance to prevent normal shaping of the body axis. However, our vital labelling experiments in embryo culture refute this hypothesis. *Lp/+* embryos undergo normal midline extension of DiO-labelled node-derived cells, with or without exposure to chlorate. In contrast, PCP-compromised *Lp/Lp* embryos show greatly reduced midline extension, confirming that defective CE is a major defect in the absence of PCP function in mice (Ybot-Gonzalez et al., 2007) as well as in other vertebrates (Wallingford et al., 2002). Furthermore, morphological features of defective CE (increased width and reduced length) are not present in chlorate-treated *Lp/+* embryos. Hence, faulty Closure 1, with subsequent development of CRN, the most severe NTD in mice and humans, is not solely the outcome of defective neuroepithelial CE, but can also arise from another embryonic abnormality in which partial loss of PCP signalling plays a non-exclusive role.

We found that inhibition of GAG sulfation changes the morphology of the Closure 1 region, prior to the normal stage of closure, suggesting that proteoglycans are involved in regulation of neuroepithelial bending. Loss of syndecan-4, an HS proteoglycan with a minority of CS chains, interacts with the *Vangl2*^*Lp*^ allele, disrupting neural tube closure in the low spine (Escobedo et al., 2013). However, Closure 1 appears normal in these mice, suggesting that the CS/HS requirement for Closure 1 may involve other proteoglycan(s). In the present study, the effect of chlorate on neural plate morphology was observed in both *+/+* and *Lp/+* embryos, arguing for a PCP-independent requirement for GAG sulfation. Chlorate-treated *+/+* individuals lose the normal ‘horseshoe’ morphology of the NE, and show delayed neural fold apposition and closure, although they are able to complete closure in most cases. By contrast, *Lp/+* embryos display more severe abnormalities of neural plate morphology, with a prominent midline bend (MHP) that is not usually seen at this axial level.

Apical constriction of neuroepithelial cells due to cytoskeletal actomyosin contraction is often viewed as the ‘principal motor’ that drives neural plate bending and closure (Sawyer et al., 2010; Suzuki et al., 2012). We recently showed that mediolateral polarization of F-actin is regulated by Vangl2. Conditional deletion of Vangl2 from neural plate and SE prevented elevation of the caudal neural folds and caused spina bifida, although Closure 1 occurred normally (Galea et al., 2018). In the present study, Vangl2 was found to be co-expressed with GAG chains in the neural plate, somitic mesoderm and SE of the Closure 1 region suggesting several potential sites of Vangl2-proteoglycan interaction. However, no obvious abnormalities were found in the intensity and localisation of actin following culture with chlorate. In particular, strong apical actin staining was present at sites of neural plate convexity in chlorate-treated *Lp/+* embryos, suggesting that deficiency of apical cytoskeletal components may not be the primary defect leading to failure of Closure 1.

A few mouse mutants have been described in which Closure 1 failure and CRN are observed, in the absence of overt PCP dysfunction. *Fgfr1* null embryos have severe defects including lack of neural tube closure and absence of somites throughout the body axis (Deng et al., 1994; Yamaguchi et al., 1994). After backcross to the C57BL/6 background, knockouts survive longer but CRN is still observed, as is failed somite formation (Hoch and Soriano, 2006). Nap1 is a regulatory component of the WAVE complex, which regulates the actin cytoskeleton and couples extracellular signals to polarized cell movement. It is expressed in both mesoderm and neural plate, with *Nap1*-deficient embryos displaying a number of developmental abnormalities including delay of mesoderm migration, absent somitogenesis and failure of Closure 1 (Rakeman and Anderson, 2006). Loss of *Oct4* gene function after E7.5 causes CRN, random heart tube orientation, failed turning, defective somitogenesis and posterior truncation (DeVeale et al., 2013). Hence, all of these gene defects cause CRN, although in each case development is severely disrupted, raising the possibility that Closure 1 failure could be a non-specific morphogenetic disruption. Nevertheless, it is striking that in each case severe abnormalities of somite formation accompany failed NT closure initiation, raising the possibility of a causal connection.

The presence of epithelial somites directly flanking the closing neural folds is a feature of Closure 1 that is not shared with neurulation at more rostral and caudal levels, where unsegmented mesoderm is adjacent to the closing NT. The level at which NT closure initiates is consistently adjacent to the early somites in mammals, as seen in mouse, human, rabbit and pig embryos (Greene et al., 1998; Peeters et al., 1998; Van Straaten et al., 2000; O’Rahilly and Müller, 2002). In chick, NT closure was found to occur by ‘buttoning’ in register with the somites, suggesting that the somites may be involved in enhancing elevation and apposition of the neural folds (Van Straaten et al., 1996). Indeed, presomitic mesoderm has been shown to compress the neural tube and notochord during axial elongation (Xiong et al., 2020). In the neurulation stage mouse embryo, CS proteoglycans are particularly abundant in the epithelial somites, and may play a mechanical role in maintaining their integrity and structure. The carboxyl and sulfate groups of GAGs trap water between their chains, generating a Donnan osmotic equilibrium that is responsible for tissue compressive stiffness, as in cartilage (Chahine et al., 2005).

In mammalian neurulation, a role for cranial mesoderm in elevation of the convex midbrain neural folds has previously been identified. Here the mesoderm is unsegmented, and the key molecular event is the synthesis of a hyaluronan-rich extracellular matrix that increases intercellular spaces and supports the elevating neural folds at the convex stage of midbrain closure (Solursh and Morriss, 1977; Morriss and Solursh, 1978). In the present study, somite morphology proved abnormal in chlorate-treated *Lp/+* embryos, consistent with a possible biomechanical role of the somites in Closure 1. Reduction of GAG sulfation led to loss of the normal ‘horse-shoe’ morphology of NE in the closure 1 region and delay of NT closure in *+/+* embryos. Added to mild CE, or other defects caused by the *Lp/+* genotype, this causes aberrant morphology of the Closure 1 region that strongly predisposes *Lp/+* embryos to failure of NT closure initiation. We propose, therefore, that the relationship between neural plate and epithelial somites may be critical for Closure 1.

In terms of human NTDs, heterozygosity for rare, non-synonymous, deleterious variants of PCP genes, including *Vangl2*, have been identified in a number of genomic studies of NTDs, including CRN (Juriloff and Harris, 2012; Robinson et al., 2012). Building on this, the present work suggests that genes encoding proteoglycan core proteins and GAG biosynthetic enzymes may also represent candidate genes that could contribute to risk of human NTDs. Array-based comparative genomic hybridization of a cohort of 189 Caucasian and Hispanic cases with non-syndromic lumbo-sacral myelomeningocele identified heterozygous deletions of glypican genes *GPC5* and *GPC6* (Bassuk et al., 2013), as a significant risk factor. Similarly, ultra-rare deleterious variants in the extracellular matrix genes *FREM2* (FRAS1 Related Extracellular Matrix 2) and *HSPG2* (perlecan) were found to be associated with myelomeningocele in a cohort of North American individuals (Au et al., 2021). Interestingly, ingestion of chlorate in drinking water has been significantly associated with risk of spina bifida in Italy (Righi et al., 2012). It will be important in future to determine whether rare variants of PCP genes, and genetic or environmental disturbance of sulfated PGs, are present in the same individuals with NTDs and therefore may contribute to NTD pathogenesis through their developmental interaction.

## MATERIALS AND METHODS

### Mouse strains and genotyping

Mouse experiments were conducted under the auspices of the UK Animals (Scientific Procedures) Act 1986 and the Medical Research Council’s Responsibility in the Use of Animals for Medical Research (1993). Inbred BALB/c mice were used for immunofluorescence and in situ hybridisation. *Vangl2*^*Lp*^ mice were maintained on the CBA/Ca background and time-mated to generate embryos with either wild-type mice within the CBA/Ca colony, or by a single generation outcross to BALB/c mice. Genotyping for the *Vangl2*^*Lp*^ allele was as described previously (Copp et al., 1994). *Vangl2*^*flox/flox*^ mice (C57BL/6J background) were a gift from Deborah Henderson (Newcastle University). *Vangl2*^*flox/flox*^ were crossed with β*-actin*^*Cre/+*^ mice (C57BL/6J background), to produce *Vangl2*^*flox/-*^; β*-actin*^*Cre/+*^. These were backcrossed to *Vangl2*^*flox/flox*^ to remove β*-actin*^*Cre/+*^, while maintaining the *Vangl2* null allele. The colony was maintained by *Vangl2*^*flox/flox*^ x *Vangl2*^*flox/-*^ matings, which were also used to generate embryos for experiments.

### Embryo culture

Embryos were explanted at E8.5 into Dulbecco’s modified Eagle’s medium containing 10% fetal bovine serum. Culture was performed in undiluted rat serum, in a roller incubator maintained at 38°C and gassed with a mixture of 5% CO2, 5% O2, 90% N2, as described (New et al., 1973; Pryor et al., 2012). For chlorate experiments, cultures were stabilized for 1 h, and then sterile aqueous sodium chlorate was added (1% volume addition) to a final concentration of 5-20 mM. The same volume of sterile distilled water was added to control cultures. Rescue experiments involved culturing embryos with 10 mM chlorate together with 10 mM exogenous sodium sulfate. GAG degrading enzymes: Chondroitinase ABC (2U), Heparitinase III (5U) or enzyme buffer, were administered by injection into the amniotic cavity of E8.5 embryos, using a hand-held, mouth-controlled glass micropipette (∼ 0.2 µl injected per embryo). Embryos were injected before the start of the culture and again after 8 h of culture. Cultures were terminated after 8 or 24 h, inspected immediately for yolk sac shape and circulation (Table S1), and presence or absence of heart beat, and then yolk sacs were removed and stored for genotyping. Embryos were scored for presence or absence of Closure 1 and somites were counted. Embryos were washed in PBS and fixed overnight in ice-cold 4% paraformaldehyde in PBS.

### Whole-mount *in situ* hybridization

*In situ* hybridization was performed as described previously (Mole et al., 2020). Primer sequences for the *Vangl2* RNA probe are 5’-GGATGCTGCTGAAAGGGAGT-3’ (forward) and 5’-GCACCGGATAGTTGGAAGGT-3’ (reverse). Sense and antisense RNA probes were transcribed from linearized plasmid DNA using a DIG RNA Labeling Kit (Roche). Sections were prepared by embedding hybridized embryos in gelatin-albumin and sectioning on a vibratome (Leica VT1000S) at 40 µm thickness.

### Section immunofluorescence

After fixation, embryos were immersed in 20% sucrose solution and placed on ice for at least 1 h. The sucrose solution was replaced with 7.5% gelatine (37 °C for 30 min) and embryos were embedded in a block of gelatine after solidification. Blocks were snap frozen in -70°C isopentane and stored at -80°C. Blocks were sectioned on a Leica cryostat at 10 μm thickness and mounted on Superfrost Plus slides (Thermo Fisher). After defrosting, slides were incubated with 200 µl of blocking buffer (10% sheep serum, 0.1% Tween in PBS) for 1 h at room temperature. For immunohistochemistry of Vangl2, blocking was in 5% normal goat serum, 0.1% Triton, 2% bovine serum albumin in PBS. The blocking solution was replaced with primary antibody (Table S2) diluted in blocking buffer and slides were kept overnight at 4 °C (200 µl per slide + parafilm). The following day, slides were washed three times in PBS and incubated in secondary antibody at 1:500 dilution in blocking buffer for 1 h at room temperature. Slides were washed three times in PBS and incubated in DAPI 1:10,000 for 3 min, washed 3 times with PBS and mounted in Mowiol. For F-actin staining, sections and whole embryos (PFA fixed only) were incubated with phalloidin (1:200, Alexa-Fluor-568–phalloidin, A12380, Life Technologies).

Chr.ABC and Hep.III specificity was confirmed on embryo cryosections, prior to enzyme usage in embryo cultures (Fig. S4). Slides were defrosted, incubated in PBS at 37 °C, and washed in PBS at room temperature. Slides were then incubated with Chr.ABC or Hep.III at 37°C for 1 h (200 µl per slide). Control slides were incubated with corresponding enzyme buffer. Slides were washed three times with PBS at room temperature, blocked with blocking buffer and then stained with corresponding primary antibodies as above.

### Whole mount immunofluorescence

After dissection, embryos were rinsed in PBS and fixed in ice-cold 4% paraformaldehyde for 1 h. Tissues were permeabilized using 0.1% Triton X-100 in PBS (PBT solution), embryos were washed for 30 min at room temperature and incubated with blocking solution (5% BSA in PBT) overnight at 4°C while rocking. Embryos were then incubated with primary antibody overnight at 4°C with rocking. Primary and secondary antibodies were diluted in blocking solution to working concentrations. Next day, embryos were washed three times in blocking solution for 1 h, rocking at room temperature, and incubated with secondary antibody for 2 h under the same conditions. Embryos were washed in blocking solution three times for 1 h rocking at room temperature, then incubated with DAPI 1:15,000 overnight rocking at 4°C. After washing three times in PBS 30 min at room temperature, embryos were stored in 0.1% sodium azide in PBS at 4°C, prior to imaging. Confocal images were obtained on a Zeiss LSM880 Observer microscope as described previously (Galea et al., 2017). In some cases (e.g. Vangl2 and E-cadherin double immunostaining, Fig. 1F), the SE and apical surface of NP were ‘isolated’ virtually using an in-house Fiji macro (IdentifyUpperSurfacev2.ijm) (Galea et al., 2018), developed to identify and extract the surface of 3D structures, available from https://github.com/DaleMoulding/Fiji-Macros.

### Assessment of axial extension by DiO injection into the embryonic node

*Vangl2*^*Lp/+*^ x *Vangl2*^*Lp/+*^ matings were performed to generate litters containing *+/+, Lp/+* and *Lp/Lp* embryos. DiO labelling of the node was performed as previously (Ybot-Gonzalez et al., 2007; Pryor et al., 2012). Some injected embryos (n = 2 per treatment/genotype group) were collected at time 0 to verify the injection site, while others were randomly distributed to either 10 mM chlorate or water control groups, and cultured for 20-24 h. After culture, embryos were assessed for health and developmental parameters, yolk sacs were stored for genotyping and fluorescence images were obtained to determine the degree of axial extension of DiO-labelled cells. Embryos were gently flattened using forceps, with ventral embryonic surface upwards, and photographed on the stage of a Leica fluorescence stereomicroscope. To confirm the presence of the dye in the midline, three cultured embryos from each treatment group were cryo-embedded for transverse sectioning. Analysis was performed blind to embryo genotype and treatment group. Distance of DiO midline extension was measured in a rostral direction from the node injection site (in the caudal region). DiO extension occurred typically as far rostrally as heart level. Embryo width was the distance between lateral edges of the somite rows, measured at the level of the 3rd/4th somites, in dorsal view. Embryo length was measured along the dorsal surface of the embryo, from forebrain to tailbud. All measurements were made in Fiji. The presence of DiO in head folds of some embryos is non-specific: caused by release of DiO into the amniotic cavity during node injection.

### Analysis of neural plate and somite morphology

Embryos were fixed in 4% PFA and exposed to CellMask™ which stains membranes non-specifically, then imaged by epifluorescence on an inverted LSM710 or LSM880 with Airyscan confocal system (Carl Zeiss Ltd, UK). For neural plate morphology, images were re-sliced in Fiji to obtain transverse sections of the Closure 1 region, located at the level of the third somite. The distance between neural folds was measured at 10 sequential positions, 20 µm apart, moving rostrocaudally along the Closure 1 region (200 µm in total). For somite morphology, the images were re-sliced in Fiji (from dorsal to ventral surface) to obtain longitudinal sections through the somites. Length (rostrocaudal orientation) and width (mediolateral orientation) measurements were taken half way though the somite at the level of third/fourth somite. Two somite pairs (4 individual somites) were measured per embryo.

### Statistical analysis

Statistical analysis of frequency of Closure 1 failure in cultured embryos was performed by Chi-squared or Fisher’s exact test. Statistical analysis of embryo length, width and midline DiO extension was performed by two-way Analysis of Variance (SPSS v.24). Analysis of neural plate morphology was performed in SPSS by mixed model analysis of distance between neural fold tips along the closure 1 region, with four fixed effects: genotype, treatment, position and somite stage. The first order interactions between fixed effects were computed per parameter. When the fixed effects were significant overall, a post-hoc Bonferroni correction was used to identify the individual sites at which the effect was significant (p-value = 0.05).

## Acknowledgements

The authors thank Dorothee Mugele for assistance with node injection and Dale Moulding for help with imaging and image analysis.

## Author contributions

Study design: ON, NDEG, PS, AJC; Methodology: ON, MM, GG, DS; Formal analysis: ON, GG, AJC; Writing - original draft: ON, AJC; Writing - review & editing: GG, MM, DS, NDEG, PS; Funding acquisition: AJC, PS.

## SUPPLEMENTARY FIGURE LEGENDS

**Figure S1**

**Co-localization of HS and CS chains with laminin.** Immunofluorescence staining on cryosections of *+/+* (A,B) and *Lp/Lp* (C,D) embryos. **(A,B)** Double immunohistochemistry: anti-laminin together with anti-CS (A, 6 somite stage) or anti-HS (B, 7 somite stage) antibodies. Laminin co-localizes with CS and HS chains at the BM of neural plate and SE at the Closure 1 site. HS staining shows an almost complete overlap with laminin whereas CS chains are also detected outside the laminin-expressing domain. (**C,D**) Both CS and HS chains localise to the basement membrane of NE and SE in the Closure 1 region of *Lp/Lp* embryos (7 somite stage), with CS chains also present in the somitic mesoderm. Distribution of GAG chains is not markedly different from *+/+* embryos, despite failed neural tube closure. Scale bars: 50 µm.

**Figure S2**

**Comparison of Vangl2 protein in wild-type and *Lp/Lp* mutant embryos. (A)** Dorsal view of the PNP region of a whole mount *+/+* embryo (14 somite stage) double immunostained for Vangl2 and E-cadherin (A-i to vi), confirming the presence of Vangl2 in SE at the posterior region. Vangl2 is expressed in broadly expressed in the neuroepithelium (A-vii to ix). **(B)** Dorsal view of caudal region of a whole mount *Lp/Lp* mutant embryo (11 somite stage) double immunostained for Vangl2 and E-cadherin. Vangl2 is absent from the mutant SE (B-i to vi), which expresses only E-cadherin. The mutant neuroepithelium is also negative for Vangl2 (B-vii to ix). Images processed by z-projection (A, B-i to vi) and single z-plane (A, B-vii to ix). Scale bars: A, B-i, ii, iii = 100 µm; A, B-iv to ix = 20 µm.

**Figure S3**

**Relationship between somite stage at start of treatment and Closure 1. (A**) Embryo somite stage in *+/+* and *Lp/+* embryos after 24 h culture with addition of either water (control) or 10 mM chlorate. Genotypes and treatments do not differ statistically in final somite stage (p = 0.448, p = 0.135 respectively). (**B**) Schematic showing somite stage(s) during the chlorate treatment period in culture, and the timing of Closure 1. Starting somite stage was calculated by subtracting the expected number of somites formed during the culture (2 h = 1 somite; 24 h = 12 somites) from the number of somites recorded at the end of the treatment. One somite was subtracted from the total somite number across all groups, to allow for the time taken to adapt to development in culture. Embryos were pooled into three ‘starting’ somite groups: 0-1, 2-3, 4-5. (**C**) Percentage of open/closed *+/+* and *Lp/+* embryos, for the calculated somite stages at the start of culture, after exposure to either water (H_2_O) or 10 mM chlorate (CL). The number of replicate embryos is indicated in each bar. Most embryos had 0-1 or 2-3 somites at the start of culture (respectively: 61% and 24% for *+/+*; 56% and 27% for *Lp/+*) and, for these pooled somite ranges, Closure 1 failed in more than 85% of chlorate-treated *Lp/+* embryos. Overall, 17% of embryos started culture in the 4-5 somite range, when it appears that chlorate-treated *Lp/+* embryos could have a lower frequency of Closure 1 failure (3/5; 60%), although numbers are too small for statistical analysis.

**Figure S4**

**Validation of GAG proteolytic enzymes and primary antibodies against CS and HS chains.** Transverse cryosections through the trunk region of wild type embryos (14-16 somite stage), pre-incubated with enzyme buffer (A,D), chondroitinase (Chr.) ABC (B,E) or heparitinase (Hep.) III (C,F), prior to immunohistochemical staining for CS or HS chains. Immunofluorescence only (i) and immunofluorescence plus DAPI images (ii) are shown. (**A-C**) CS chains show the distinct basement membrane pattern of staining in buffer-treated controls (A) whereas this is abolished by Chr.ABC pre-treatment (B). Hep.III pre-treatment has minimal effect on CS staining (C). (**D-F**) HS chains are present in basement membranes of SE (strong) and neuroepithelium (weaker) after buffer pre-treatment (D). Hep.III-treated sections show strong reduction of HS staining (F), whereas Chr.ABC does not affect the expression pattern of HS chains (E). Scale bars: 50 µm.

**Figure S5**

**Quantification of chlorate effects on neural plate morphology at the Closure 1 site.** *+/+* and *Lp/+* embryos were cultured for 8 h with addition of 10 mM chlorate or water, fixed, stained with CellMask™ and imaged using confocal microscopy for morphological analysis. **(A)** Dorsal confocal view (rostral to left) showing the converging neural folds at the Closure 1 site of a normally developing embryo. **(B)** Images were re-sliced in Fiji to obtain transverse sections of the Closure 1 region, located at the level of the third somite (along dashed line in A). The distance between the neural fold tips (‘D’ in B) was measured at 10 sequential positions with 20 µm spacing, moving rostrocaudally along the body axis and centred upon the Closure 1 region. **(C,D)** Distance between neural fold tips averaged across all positions (C) and at individual positions (D) at the 5, 6 and 7 somite stages. Distance between neural fold tips decreases most rapidly with increasing somite stage in *+/+* control embryos (black bars, lines), with closure essentially complete at 7 somites. Chlorate-treated *+/+* (grey bars, lines) and water-treated *Lp/+* (green bars, lines) embryos show progressive but delayed closure. Chlorate-treated *Lp/+* embryos (orange bars, lines) show little progression towards closure. Data values: mean +/-SEM, with a minimum of 4 embryos per genotype/treatment group. * p < 0.05; **p < 0.01;***p < 0.001. Scale bar: A,B = 50 µm.

## SUPPLEMENTARY TABLES

**Table S1.**
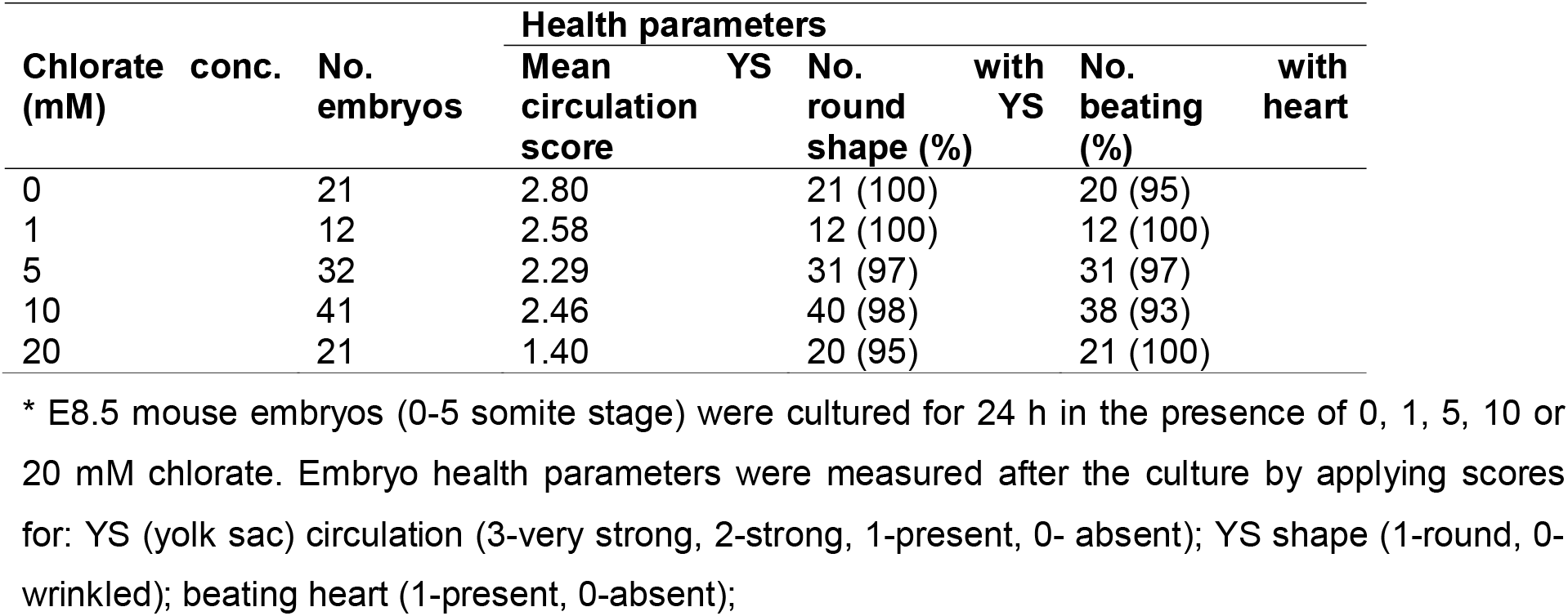
Effect of increasing chlorate concentration on embryo health parameters during culture *.

**Table S2.**
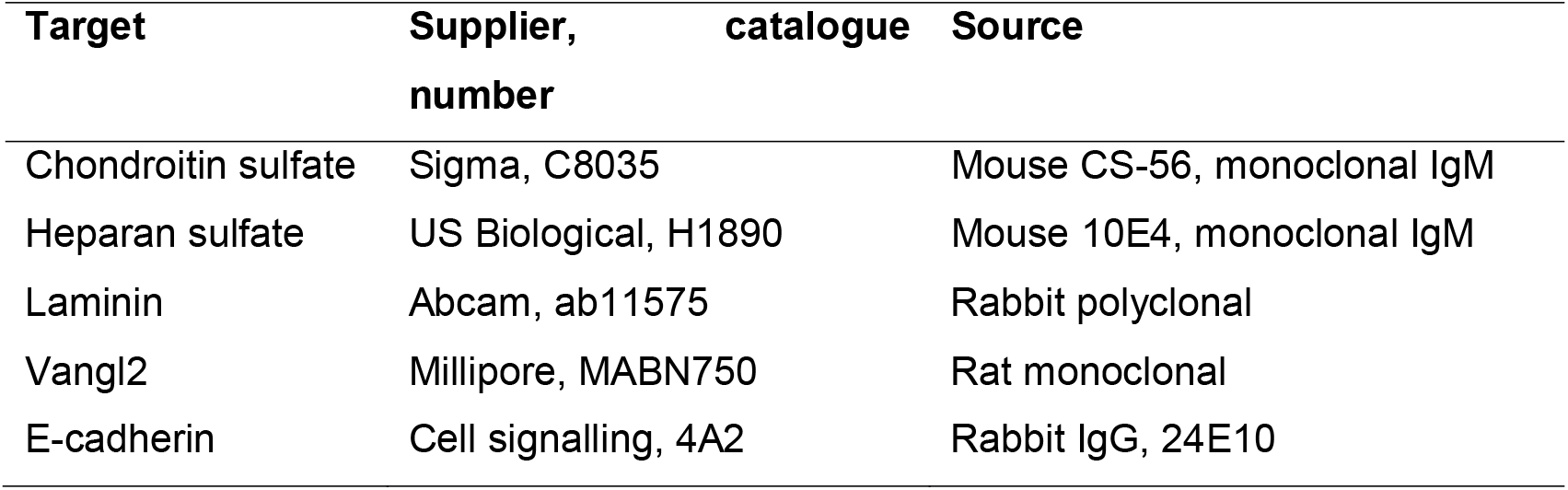
Primary antibodies used in the study.

